# Identification of genetic variants regulating the abundance of clinically relevant plasma proteins using the Diversity Outbred mouse model

**DOI:** 10.1101/2020.11.04.367938

**Authors:** Stéphanie Philtjens, Dominic J. Acri, Byungwook Kim, Hyewon Kim, Jungsu Kim

**Author notes:** Corresponding author: (JK). These authors contributed equally to this work (Stéphanie Philtjens, Dominic J. Acri).

## Abstract

Although there have been numerous expression quantitative trait loci (eQTL) studies, the effect of genetic variants on the levels of multiple plasma proteins still warrants more systematic investigation. To identify genetic modifiers that influence the levels of clinically relevant plasma proteins, we performed protein quantitative trait locus (pQTL) mapping on 92 proteins using the Diversity Outbred (DO) mouse population and identified 12 significant *cis* and 6 *trans* pQTL. Among them, we discovered coding variants in a *cis*-pQTL in *Ahr* and a *trans-*pQTL in *Rfx1* for the IL-17A protein. Our study reports an innovative pipeline for the identification of genetic modifiers that may be targeted for drug development.

**Author Summary:** Blood plasma is a body fluid that can be collected in a noninvasive way to detect diseases, such as autoimmune disease. However, it is known that plasma protein levels are affected by both the environment and genetic background. To determine the effect of genetics on plasma protein levels in human, one needs a rather large sample size. To overcome this critical issue, a mouse model, the Diversity Outbred (DO), was established that is genetically as diverse as the human population. In this study, we used N=140 DO mice and genotyped over 140,000 variants. In addition, we measured the levels of 92 proteins in plasma of these DO mice using Olink Proteomics technology. The proteins detected in this panel are known to be detectable in human plasma, making our study translatable to human. We identified 18 significant protein quantitative trait loci. Furthermore, we describe an analysis pipeline that allows for the detection of a single gene in the locus that is responsible for the differences in protein levels. We identified how variants in the Regulatory Factor X1 (Rfx1) gene regulates Interleukin-17A (IL-17A) plasma levels. Our study reports an innovative approach to identify genetic modifiers that may be targeted for drug development.

## Introduction

Proteins expressed in blood plasma are diverse and their levels are dependent on environmental factors and genetic background [1]. Genes that influence the expression of other genes are called modifiers and can be detected using quantitative trait loci (QTL) mapping [2, 3]. Although protein QTLs (pQTLs) have been detected in the plasma of humans and mice, they mainly detected *cis*-acting pQTLs, not *trans-*acting pQTLs, due to a lack of statistical power, a low genetic diversity, or low-throughput protein level screening platforms [1, 4–9].

To overcome the lack of genetic diversity in mouse models, the Diversity Outbred (DO) mouse model was established through a multiparent paradigm from eight founder strains, comprising five inbred and three wild-derived strains [10, 11]. The combination of eight founder strains results in a much greater level of genetic diversity than existing recombinant inbred lines and the genetic variants are more uniformly distributed across the genome than in other genetic reference populations [12]. Each DO mouse is a genetically unique individual with a high level of allelic heterozygosity, providing precision for mapping QTL with relatively small sample sizes compared to human mapping studies. For example, QTL mapping in these multiparent populations resulted in the identification of genetic modifiers for the viral response [13, 14], kidney disease [15], atherosclerosis [16] and heart size [17].

We measured the abundance of clinically relevant plasma proteins in 140 DO mice using the Olink Mouse Exploratory Panel. This proximity by extension assay (PEA) measures 92 proteins in just 1 μL of plasma [9, 18]. In addition, each DO mouse was genotyped using the third generation Mouse Universal Genotyping Assay (MUGA), the GigaMUGA [19]. To identify new modifier genes that influence the level of plasma proteins, we performed pQTL mapping (Fig 1a). To the best of our knowledge, this is the first discovery study to report *trans-*acting pQTLs in the plasma of DO mice.

**Fig 1.**
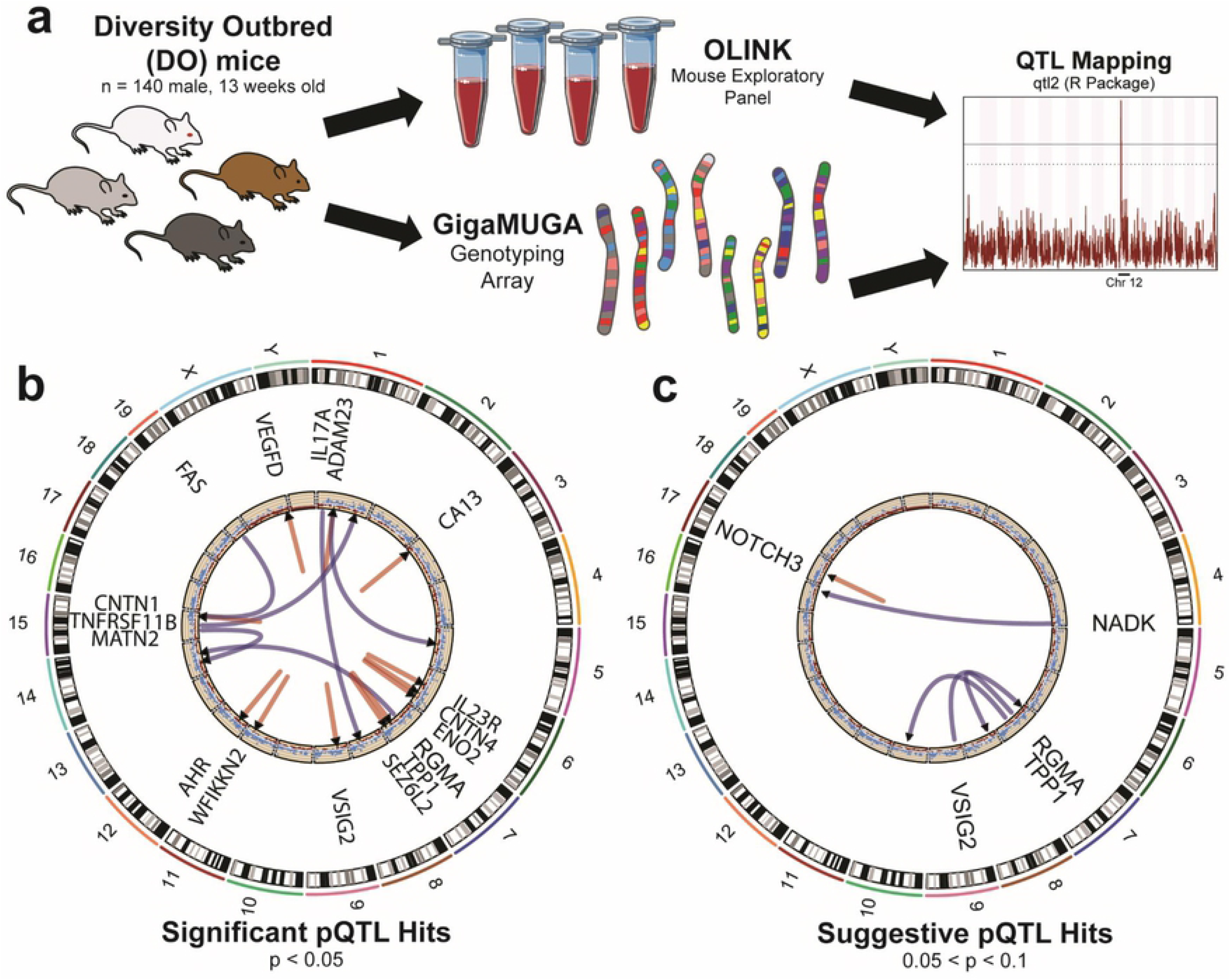
pQTL mapping of plasma proteins in the DO mouse model reveals 18 significant and 5 suggestive candidate modifier loci. **(a)** Workflow for pQTL mapping. **(b)** pQTL mapping of plasma proteins detected by the Olink Mouse Exploratory panel identified 18 significant (P-value < 0.05) pQTL. The first track includes the protein symbols mapped to their respective loci in the mouse genome (mm10). The second track is a scatterplot showing the density of the GigaMUGA genotyping array (# markers/0.5Mb; red: ≤ 20 markers; light blue: ≤ 30 markers; dark blue: ≥ 30 markers; clipped at 100 markers/0.5Mb). The inner track displays *trans* (blue) and *cis* (red) pQTL with a black triangle at the associated locus. **(c)** pQTL mapping of plasma proteins detected 5 suggestive pQTL (0.05 < P-value < 0.1). Visualization is similar to above, with *trans* (blue) and *cis* (red) pQTL.

## Results and Discussion

To identify modifier genes that affect levels of plasma proteins, we performed pQTL mapping for the analytes measured using the Mouse Exploratory Panel from Olink Proteomics with genetic relatedness as a covariate. Significant pQTL were defined by a genome-wide P-value < 0.05, while suggestive pQTL were defined by 0.05 < P-value < 0.1. We identified a total of 18 significant and 5 suggestive pQTL (Fig 1b and c, Table 1). Six of the 18 significant pQTL were *trans* pQTL (Fig 1b, blue lines) and 12 were *cis* pQTL (Fig 1b, red lines), while four of the suggestive pQTL were *trans* pQTL (Fig 1c, blue lines) and one was a *cis* pQTL (Fig 1c, red lines). A list of all identified pQTL can be found in Table 1.

**Table 1.** pQTL for plasma proteins in DO mice.

| Protein | Chromosome<br>with pQTL<br>peak | Peak<br>LOD<sup>1</sup> | Position<br>(Interval, cM) | Cis/trans | Significant<br>($p < 0.05$)<sup>2</sup> |
| --- | --- | --- | --- | --- | --- |
| ADAM23 | 1 | 10.2 | 28.7 (28.4-28.8) | *cis* | Yes |
| ADAM23 | 5 | 7.5 | 27.8 (27.4-28.9) | *trans* | No |
| AHR | 12 | 12.8 | 14.5 (14.2-16.2) | *cis* | Yes |
| CA13 | 3 | 22.9 | 1.9 (1.8-2.3) | *cis* | Yes |
| CNTN1 | 15 | 21.7 | 40.8 (40.7-40.8) | *cis* | Yes |
| CNTN4 | 6 | 37.4 | 45 (44.9-45) | *cis* | Yes |
| ENO2 | 6 | 21.4 | 55.5 (55.5-55.6) | *cis* | Yes |
| FAS | 15 | 8.3 | 44.8 (44.5-45.5) | *trans* | Yes |
| IL17A | 8 | 8.3 | 35.5 (35.4-36.3) | *trans* | Yes |
| IL23R | 6 | 35.8 | 28.1 (27-28.1) | *cis* | Yes |
| MATN2 | 14 | 8.7 | 16.7 (16.3-16.8) | *trans* | Yes |
| NADK | 16 | 7.4 | 34 (29.7-34.9) | *trans* | No |
| NOTCH3 | 17 | 7.5 | 15.9 (14.6-16.4) | *cis* | No |
| RGMA | 8 | 14.1 | 18.8 (18.8-18.8) | *trans* | No |
| RGMA | 14 | 16.1 | 35 (35-35) | *trans* | Yes |
| SEZ6L2 | 7 | 29.6 | 60 (60-60.2) | *cis* | Yes |
| TNFRSF11B | 1 | 8.4 | 60.3 (60.2-61) | *trans* | Yes |
| TPP1 | 7 | 8.8 | 45.8 (45.5-47) | *cis* | Yes |
| TPP1 | 10 | 7.7 | 37.7 (37.4-39.3) | *trans* | No |
| VEGFD | X | 12.1 | 68.7 (68.3-68.8) | *cis* | Yes |
| VSIG2 | 7 | 7.4 | 8 (7.8-27) | *trans* | No |
| VSIG2 | 9 | 10.1 | 15.5 (15.5-17.3) | *cis* | Yes |
| WFIKKN2 | 11 | 10 | 51.5 (50.6-51.9) | *cis* | Yes |

<sup>1</sup>LOD = logarithm of the odds; <sup>2</sup>“Yes” indicates a $p < 0.05$ while “No” corresponds to $0.1 < p < 0.05$ .

We identified a significant *cis* pQTL for the aryl hydrocarbon receptor (AHR) protein with a logarithm of the odds (LOD) score of 12.8 and a peak located at 33.3 Mb on chromosome 12 (Fig 1b and 2a). Fine mapping of this pQTL resulted in the identification of the missense variant p.K432R in *Ahr*, where the presence of the minor G allele significantly increased the protein abundance of AHR (P-value = 3.41 × 10^−7^, Fig 2b). Protein sequence alignment showed conservation of the lysine at position 432 between mouse and human (S1a Fig). In addition, we identified a variant located in the 5’ untranslated region (UTR) of *Ahr*, where a significant decrease in AHR protein levels was observed with the presence of the minor A allele of c.-100G>A (P-value = 6.3 × 10-9, Fig 2b). Although p.K432R is not located in a known protein domain [20] nor is c.-100G>A located in a predicted 5’ UTR motif, our data do suggest that coding variants in *Ahr* are responsible for the differences in AHR protein abundance.

**Fig 2.**
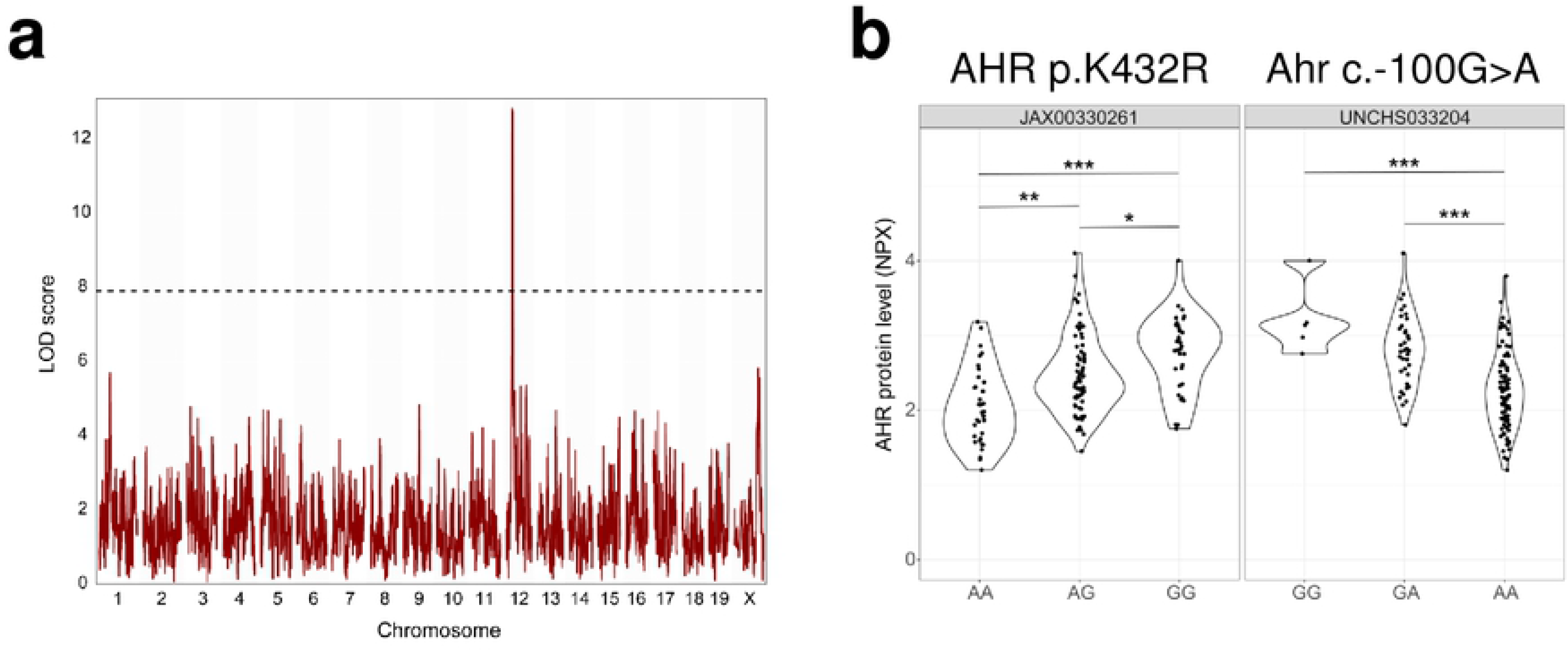
Variants in Ahr have a significant effect on AHR levels in plasma of DO mice. **(a)** Genome scan of the AHR protein shows a significant peak on chromosome 12. Chromosome position is on the x-axis and LOD score on the y-axis. Dashed line at P-value ≤ 0.05 significance threshold. **(b)** Violin plots showing the effect of the p.K432R and c.-100G>A genotypes on AHR protein abundance. *** P-value < 0.001; ** P-value < 0.01; * P-value < 0.05

One of the significant *trans* pQTL we identified was for the interleukin 17A (IL-17A) protein with a LOD score of 8.3 and a peak at 84.8 Mb on chromosome 8 (Fig 3a). Fine mapping of this region resulted in a locus of approximately 3 Mb in size (Fig 3b) [21, 22]. To identify the gene that could explain the variation in IL-17A protein levels, we investigated the 163 genes located in a 4 Mb interval around the peak. The “Shortest Path” algorithm from the MetaCoreTM “Build Network” function was used to connect the candidate modifier genes with IL-17A. A direct interaction was identified between regulatory factor X1 (RFX1) and IL-17A (Fig 3c, green highlight). Therefore, we hypothesize that *Rfx1* is a genetic modifier for IL-17A abundance in plasma. We identified two coding variants, p.F612S and p.A724, (Fig 3d) in *Rfx1*, as well as one in the 3’ UTR (c.*935G>A, Fig 3d). Although the presence of one minor C allele of p.F612S did not affect IL-17A protein levels, we did observe a significant decrease in IL-17A protein levels in DO mice carrying two minor C alleles (P-value = 0.0036, Fig 3d) compared to one. For the silent variant, p.A724, we did not observe any significant effect on the protein abundance of IL-17A (Fig 3d), as might be expected. Because p.F612 is conserved between mouse and human (S1b Fig), this amino acid residue might regulate the function of RFX1 protein. Based on the finding by Zhao et al. [23], it is tempting to hypothesize that the homozygous p.F612S increases the stability or activity of RFX1, leading to a decrease in IL-17A protein abundance. However, further research is warranted to test this hypothesis, which is outside the scope of this pQTL study. We also observed a significant decrease in IL-17A protein abundance in heterozygous carriers of c.*935G>A compared to homozygous wild type carriers (P-value = 0.0049, Fig 3d), while no significant difference was observed in homozygous carriers of the minor A allele. The c.*935G>A variant was also conserved between human and mouse (S1c Fig). Because microRNAs are known to bind 3’ UTR and downregulate gene expression, we searched for potential microRNAs that might bind around this SNP. However, this variant is not located in a known miRNA binding site according to PicTar and miRDB database [24, 25]. Our data suggest that *RFX1* is a genetic modifier for IL-17A plasma levels. This hypothesis is further supported by a recent article by Zhao and colleagues [23]. They showed that the levels of IL-17A mRNA and protein significantly increased after knock-down of RFX1 in CD4+ T cells, while a decrease in IL-17A level was observed after overexpressing RFX1. Furthermore, they demonstrated that RFX1 directly affects IL-17A expression by binding one of its two binding sites upstream of the transcription start site of the *IL17A* gene. RFX1 also played an important role in methylation and acetylation of the promoter region of *IL17A* [23]. The results from Zhao et al. [23] provide functional evidence directly linking RFX1 and IL-17A. This proof-of-concept case study clearly demonstrates the high potential of our approach to the identification of genetic modifiers for many other proteins.

**Fig 3.**
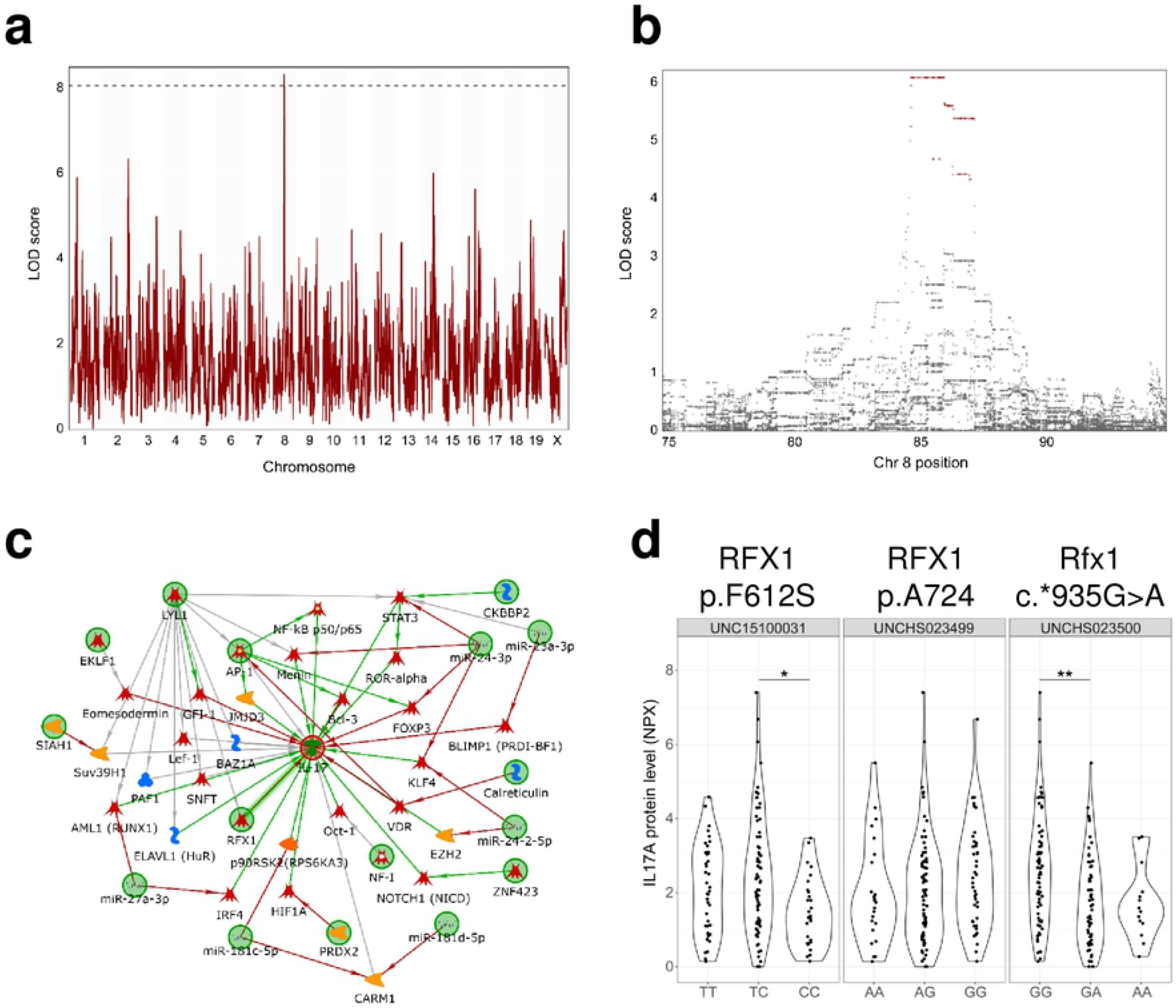
Variants in *Rfx1* have a significant effect on IL-17A levels in plasma of DO mice. **(a)** Genome scan for the IL-17A protein shows a significant peak on chromosome 8. **(b)** Association mapping of the IL-17A pQTL. Zoom in on chromosome 8 showing the QTL support interval. Magenta marks SNPs with a LOD drop < 2 from the top SNP. **(c)** MetaCore™ network analysis connecting the genes located in the pQTL locus on chromosome 8 for IL-17A. Green circles indicate either IL-17A or genes that were present in the locus. **(d)** Violin plots showing the effect of the RFX1 p.F612S, p.A724 and c.*935G>A genotypes on IL-17A protein abundance. ** P-value < 0.01; * P-value < 0.05

In summary, we identified 18 significant pQTLs for the discovery of novel genetic modifiers that control levels of plasma proteins. Of note, we found new coding variants in *Ahr* that alter AHR protein abundance in plasma. We also report both coding and non-coding variants in the *Rfx1* gene that alter IL-17A plasma protein abundance, replicating a known connection between RFX1 and IL-17A in mouse and human [23]. Additionally, we show that the genetic diversity of the DO mouse model makes it possible to identify *trans* pQTL that can easily be translated to human, making this mouse model suitable for translational discovery studies. Importantly, we demonstrated that fewer than 150 DO mice are sufficient to identify statistically significant pQTL compared to the larger number of DO mice used in previous studies [16, 17, 26–30]. The methodology detailed in this study can be used to unravel the complex mechanisms between genetic loci and protein abundance.

## Materials and Methods

### Ethics statement

All animal experiments were performed in accordance with the approved animal protocol and guidelines established by the Institutional Animal Car Committee at Mayo Clinic, Jacksonville, FL (#A00003398-00).

### Mice

Male DO mice (n=140, Stock No. 009376) were obtained in two batches from The Jackson Laboratory at 4-weeks of age and at generation 28 (G28 litter 1 and G28 litter 2) of outcrossing. The mice were housed under standard laboratory conditions on 12-h light:dark cycles in a specific pathogen-free environment at Mayo Clinic Jacksonville, Florida. At 13-weeks of age, the mice were anesthetized using ketamine (90 mg/kg, *i.p.*) and xylazine (10 mg/kg, *i.p.*). Around 200 μL of whole blood was collected into an EDTA coated tube (BrainTree Scientific Inc, Cat. No. MV-CB300-16444-BX) from the left ventricle and centrifuged at 2,000 x *g* for 15 minutes to separate out plasma.

### Genotyping

Tail samples were collected and sent to GeneSeek (Neogen) for genotyping on the GigaMUGA. The GigaMUGA contains 143,259 genetic markers that were specifically designed for genetic mapping in the DO mouse population [19, 31]. Genotype quality was assessed using the R package argyle [32].

### Olink Mouse Exploratory Panel

Plasma samples were randomly distributed across 96-well plates and shipped to Olink Proteomics (Olink Proteomics, Uppsala, Sweden). The Olink technology allows the measurement of up to 92 proteins in 1 μL of plasma using the proximity extension assay (PEA). Distinct polyclonal oligonucleotide-labeled antibodies were added to the samples and bind their target proteins. Proteins were quantified by real-time quantitative PCR of the amplified oligonucleotide tags [33]. Primary data acquisition and quality control analysis were performed by Olink Proteomics that produced the Normalized Protein eXpression (NPX), an arbitrary unit used by Olink in Log2 scale [9, 33]. 83% (76/92) of the detected proteins were detected in more than 10% of the samples and were used for pQTL mapping (S1 Table and S1 File).

### pQTL mapping

pQTL mapping was performed using the R package R/qtl2 [34]. Founder haplotype probabilities were predicted using a Hidden Markov Model adapted for multi-parent populations and the protein abundance was regressed on these founder haplotype probabilities. To account for genetic similarity between mice, the kinship matrix was determined based on the leave-one-chromosome-out method. Genome scans were performed and a random effect was included to account for kinship. Significance thresholds were determined using 1,000 permutations and the mapping statistic is the logarithm of the odds (LOD) score. pQTL significance intervals were defined by the 95% Bayesian credible interval. Significance threshold of a pQTL was set at P<0.05, and suggestive at 0.05 <P< 0.1. Genes within 2 Mb of the top SNPs, both upstream and downstream, were considered as candidate modifiers for further study.

### Motif and miRNA binding site discovery, sequence alignment and network analysis

To identify whether the 5’ UTR variant in Ahr (c.-100G>A) was located in an UTR motif, the online tools MEME Suite 5.1.1 [35] (http://meme-suite.org/index.html, Accessed on 05/26/2020) and Regma 2.0 [36] (http://regma2.mbc.nctu.edu.tw/index.html, Accessed on 05/26/2020) were used. To identify whether the 3’ UTR variant in Rfx1 (c.*935G>A) was located in a miRNA binding site, the online tools PicTar [25] (https://pictar.mdc-berlin.de/, Accessed on 05/26/2020) and miRDB [24] (http://mirdb.org/, Accessed on 05/26/2020) were used. The Basic Local Alignment Search Tool (BLAST) was used to compare mouse and human sequences at the level of the transcript and protein. Coding variants were numbered relative to the translation initiation codon in the Ahr transcript (RefSeq NM_013464.4) or Rfx1 transcript (mouse: RefSeq NM_009055.4; human: RefSeq NM_002918). Amino acid numbering is according to AHR (human: GenPept accession number NP_001612.1; mouse: GenPept accession number NP_038492.1) or RFX1 (human: GenPept accession number NP_002909.4; mouse: GenPept accession number NP_033081.3). An overview of all identified variants and their position can be found in the S3 Table. Network analysis was performed using the “Build Network” function in MetaCoreTM (Clarivate Analytics, Accessed on 03/25/2020). The “Shortest path” algorithm was used to connect the genes located in the associated locus (“from”) with IL17A (“to”). The maximum number of steps allowed between the candidate modifier genes and IL17A was two, the minimum number of steps allowed using the “Shortest path”.

### Statistical analysis

One-way ANOVA and Tukey multiple pairwise-comparison analyses were performed using R (R version 3.5.2).

## Acknowledgments

We thank Matthew Heck for his contribution in dissecting the DO mice. We would also like to thank Dr. Luke Dabin for his critical revision of this manuscript. Illustrations in this paper were created using Seriver Medical Art templates, which are licensed under a Creative Commons Attribution 3.0 Unported License.

## Availability of data and materials

The Olink Proteomics data are included as supplementary data, while the genotype data of the DO mice are available from the corresponding author.

## Authors’ contributions

Conceptualization: Stéphanie Philtjens, Dominic James Acri, Jungsu Kim

Data Curation: Stéphanie Philtjens

Formal Analysis: Stéphanie Philtjens, Dominic James Acri

Funding Acquisition: Jungsu Kim

Investigation: Stéphanie Philtjens, Dominic James Acri, Jungsu Kim

Methodology: Stéphanie Philtjens, Dominic James Acri, Jungsu Kim

Project administration: Stéphanie Philtjens, Jungsu Kim

Resources: Stéphanie Philtjens, Byungwook Kim, Hyewon Kim, Jungsu Kim

Supervision: Jungsu Kim

Validation: Stéphanie Philtjens, Jungsu Kim

Visualization: Stéphanie Philtjens, Jungsu Kim

Writing – Original Draft Preparation: Stéphanie Philtjens, Dominic James Acri, Jungsu Kim

Writing – Review & Editing: Stéphanie Philtjens, Dominic James Acri, Byungwook Kim, Jungsu Kim

## Supporting information

**S1 Fig. The coding variants in *Ahr* and *Rfx1* are conserved between mouse and human. (a)** Amino acid alignment for the p.K438R variant in *Ahr* is shown for mouse (NP_038492.1) and human (NP_001612.1). **(b)** Amino acid alignment for the p.F612S variant in *Rfx1* is shown for mouse (NP_033081.3) and human (NP_002909.4). **(c)** Base pair alignment for the 3’ UTR variant c.*935G>A in *Rfx1* is shown for mouse (NM_009055.4) and human (NM_002918).

**S1 Table. Overview of proteins analyzed using the Olink Mouse Exploratory Panel**

**S2 Table. List of identified variants.** ^1^Relative to the translation initiation codon in NM_013464.4 for Ahr and NM_009055.4 for Rfx1; ^2^According to NP_038492.1 for AHR and NP_033081.3 for RFX1; ^3^SNP identifier as used by GigaMUGA (GeneSeek, Neogen).

**S1 File. Olink Mouse Exploratory panel protein abundane measurements** (XLSX)

